# deepMc: deep Matrix Completion for imputation of single cell RNA-seq data

**DOI:** 10.1101/387621

**Authors:** Aanchal Mongia, Debarka Sengupta, Angshul Majumdar

## Abstract

Single cell RNA-seq has fueled discovery and innovation in medicine over the past few years and is useful for studying cellular responses at individual cell resolution. But, due to paucity of starting RNA, the data acquired is highly sparse. To address this, We propose a deep matrix factorization based method, deepMc, to impute missing values in gene-expression data. For the deep architecture of our approach, We draw our motivation from great success of deep learning in solving various Machine learning problems. In this work, We support our method with positive results on several evaluation metrics like clustering of cell populations, differential expression analysis and cell type separability.

## 1 Introduction

Bulk RNA sequencing has traditionally been used for parallel screening of thousands of genes by revealing a global view of averaged expression levels. Single cell RNA sequencing (scRNA-seq), on the contrary, enables transcriptomic analysis and measurement of gene expressions at the single cell level, thus providing more perceptivity into functioning of individual cells. Over the past few years, scRNA-seq has transformed the field of functional biology and genomics [1] by enabling characterization of phenotypic diversity among seemingly similar cells [2, 3, 4]. This unique feature has been proved critical in characterizing cancer heterogeneity [5, 6], identification of new rare cell types and understanding the dynamics of transcriptional changes during development [7, 8, 9].

However, this powerful technology suffers from a number of sources of biological and technical noise, the major one being lack of starting mRNA captured in individual cells. Due to small quantities transcripts are frequently missed during the reverse transcription step. This leads to ‘dropout’ events, where only a fraction of transcriptome of each cell is detected during the sequencing step [10], leading to a sparse gene-expression matrix. This often happens with the lowly expressed genes. Excluding these genes from analysis may not be the most viable solution as many of the transcription factors and cell surface markers are sacrificed in this process [11]. Also, variability in dropout rate across individual cells or cell types, works as a confounding factor for a number of downstream analyses [12, 13]. It has been shown for scRNA-seq datasets that the first principal components highly correlate with proportion of dropouts across individual transcriptomes. So, efficient imputation strategies need to be devised to recover the lost gene-expression for more accurate gene expression measurements in scRNA-seq datasets.

To our knowledge, Recent efforts to cater this problem include MAGIC [11], scImpute [14] and drImpute [15]. MAGIC is based on the idea of heat diffusion and uses a neighborhood based heuristic to infer the missing values. It imputes missing values by sharing information across similar cells. ScImpute, first learns each gene’s dropout probability in each cell based on a mixture of Gamma and Normal distributions. It then imputes the dropout values in a cell by borrowing information of the same gene in other similar cells, which are selected based on the genes unlikely affected by dropout events. ScImpute claims to have better performance than MAGIC. Since the sources of technical noise and biases are not known [13], parametric modeling of single cell expression is challenging. Moreover, there is clear lack of consensus about the choice of probability density function. Another proposed imputation method, Drimpute, repeatedly identifies similar cells based on clustering, and performs imputation multiple times by averaging the expression values from similar cells, followed by averaging multiple estimations for final imputation.

We propose deepMc, a deep Matrix Factorization based imputation technique for scRNA-seq data. Our technique does not assume any distribution for gene expression, outperforms other proposed imputation techniques in most experimental conditions and scales gracefully for a large droplet-sequencing data containing transcriptomes in the order of thousands like PBMCs, having 68K cells (on which other imputation strategies could not run). We believe that superior performance, combined with scalability will make deepMc the method of choice for imputing scRNA-seq data.

## 2 Methods

### 2.1 Dataset description

We used scRNA-seq datasets from four different studies for performing various experiments.

- **PBMC data:** This scRNA-seq data features fresh ∼68,000 PBMCs (peripheral blood mononuclear cells), collected from a healthy donor. Single cell expression profiles of 11 purified subpopulations of PBMCs are used as reference for cell type annotation [16]. We downloaded the count data from 10x Genomics website.
- **Jurkat-293T data:** This dataset contains expression profiles of Jurkat and 293T cells, mixed *in vitro* at equal proportions (50:50). All ∼ 3,300 cells of this data are annotated based on the expressions of cell-type specific markers [16]. Cells expressing CD3D are assigned Jurkat, while those expressing XIST are assigned 293T. This dataset is also available at 10x Genomics website.
- **Preimplantation data:** This is an scRNA-seq data of mouse preimplantation embryos. It contains expression profiles of ∼ 300 cells from zygote, early 2-cell stage, middle 2-cell stage, late 2-cell stage, 4-cell stage, 8-cell stage, 16-cell stage, early blastocyst, middle blastocyst and late blastocyst stages. The first generation of mouse strain crosses were used for studying monoallelic expression. We downloaded the count data from Gene Expression Omnibus (GSE45719) [7].

### 2.2 Data preprocessing

Steps involved in preprocessing of raw scRNA-seq data are enumerated below.

- **Data filtering:** Cell filtering was not necessary for Jurkat-293T and PBMC data as 10x genomics default pipeline takes care of that. Preimplantation data had small number of cells, of which no cell was found with alarmingly poor total read count (minimum total read count being 2,56,681. If a gene was detected with *≥* 3 reads in at least 3 cells we considered it expressed. We ignored the remaining genes.
- **Library-size Normalization:** Expression matrices were normalized by first dividing each read count by the total counts in each cell, and then by multiplying with the median of the total read counts across cells.
- **Gene Selection:** For each expression data top 1000 high-dispersion (coefficient of variance) genes were kept for imputation and further analyses.
- **Log Normalization:** A copy of the matrices were log_2_ transformed following addition of 1 as pseudocount.
- **Imputation:** For various experiments, log transformed expression matrix was used as input for imputation.

### 2.3 Brief Literature Review

In scRNA-seq data, only a fraction of transcriptome of each cell is captured due to insufficient input RNA. This makes the measured gene expression a partially observed version of the actual data with no dropout events. Our aim is to impute the droputs or missing values. The formal model for measurement can be expressed as follows:

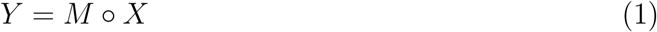

where, ◦ is the hadamard product and M is the binary mask containing 1’s where Y contains an entry and 0’s elsewhere. Here, X represent the count matrix with no dropouts, that needs to be estimated.

This is an under-determined linear inverse problem and hence has infinitely many solutions; therefore there are multiple ways to approach it. The traditional approach is based on matrix factorization [17]. In the simplest form, it assumes that X is a low rank matrix and hence can be expressed as a product of one thin (U) and one fat (V) matrix: X=UV. Incorporating this model into (1) leads to the standard matrix factorization problem.

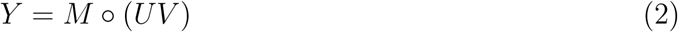

This is usually solved by alternating least squares (ALS) technique to recover *U* and *V*.

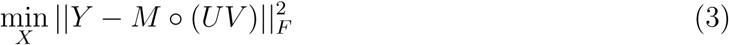

Even though more sophisticated solutions exists, for most practical problems the simple ALS is good enough.

Although matrix factorization techniques have been well known since the 90’s [18, 19], it suffers from certain fundamental problems. The factorization approach (3) is bi-linear and hence non-convex, and therefore suffers from non-convergence and non-uniqueness. A mathematically better, albeit more abstract approach to solve (1) is to directly solve for a low-rank X from (1). This is achieved via nuclear norm minimization [20, 21].

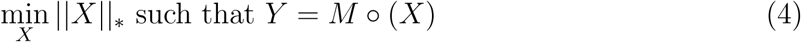

The nuclear norm *||X||*_*∗*_ is the closest convex surrogate for rank of a matrix. Solving the exact rank minimization problem is to be NP hard, but (4) can be solved via semi-definite programming (today more efficient techniques exist). Such a nuclear norm minimization technique has been used for estimating missing values in bulk data [22].

### 2.4 Overview of deepMc/ Proposed approach

In recent times, deep learning has permeated almost every aspect of computational science. Bioinformatics is not an exception. Although slightly dated [23] is a comprehensive treatise into the early applications of deep learning in this area. Our current work is motivated by the success of deep matrix factorization [24, 25] and deep dictionary learning [26]. The basic idea in there is to factor the data matrix into several layers of basis and a final layer of coefficients; shown here for three levels

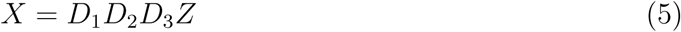

For visual understanding, please see deepMc architecture shown in Figure 1.

**Figure 1:**
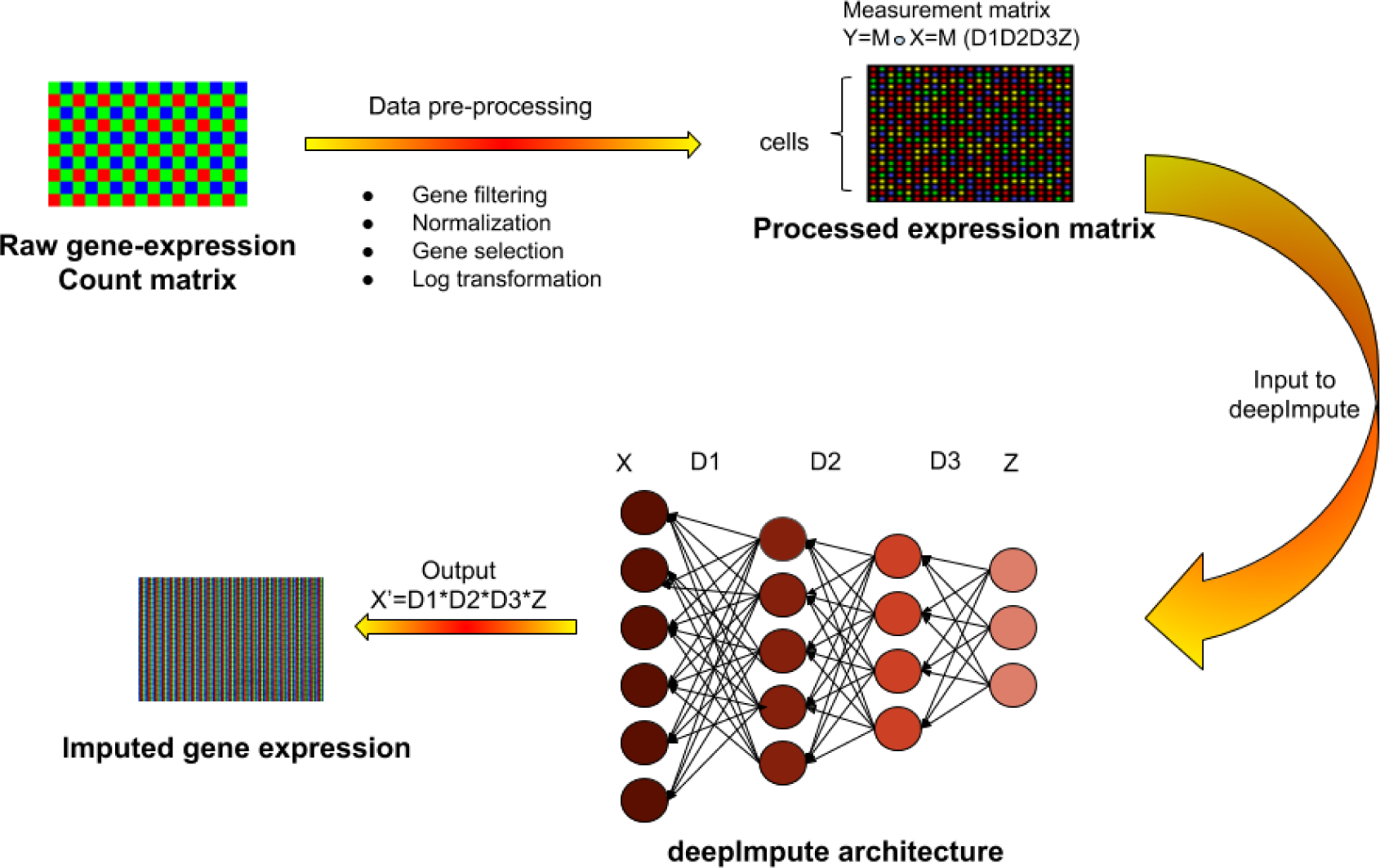
Overview of deepMc pipeline for imputing single cell RNA sequencing data. Raw read counts were filtered for significantly expressed genes and then normalized by Library size. Then, only the top 1000 genes with highest dispersion were kept and the expression data was Log2 transformed (after adding a pseudo count of 1). This pre-processed expression matrix (Y) is treated as the measurement/observation matrix (and fed as input to deepMc) from which the gene expressions of the complete matrix (X) need to be recovered by learning several layers of basis D1, D2, D3 and a final layer of coefficients Z.

Note that this is a feedbackward neural network, the connections are from the nodes towards the input. This is because matrix factorization is a synthesis formulation. Incorporating the deep matrix factorization formulation into (1) leads to,

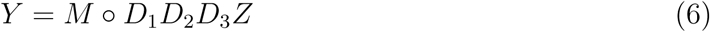

Our task is to solve the different layers of basis (D1, D2, D3) and the coefficients (Z) by solving the least squares objective function.

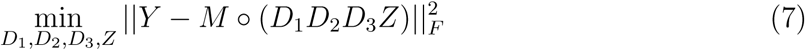

Note that we cannot use techniques derived in [24, 25] for our purpose; this is because they operated on the full data where as we need to derive for partially observed data. The derivation for (7) is relegated to the supplementary.

## 3 Results

### 3.1 Contribution

The major contribution of this work in bio-informatics is that, we show how abstract albeit powerful mathematical models can yield better results compared to heuristic techniques based on biological understanding such as MAGIC [27], drImpute [15] and scImpute [28]. This is the first work that models imputation of single cell RNA sequences as a matrix completion problem. Without any biological prior, our method outperforms the aforesaid techniques.

The other contribution of this work extends beyond bioinformatics, and is a fundamental contribution to machine learning/signal processing. This is the first work that shows how matrix completion can be framed as a deep learning problem. The proposed technique will impact other areas of applied matrix completion, viz. collaborative filtering [29], system identification [30], direction of arrival estimation [31] etc.

Also, our proposed method gracefully scales for large datasets, which other imputation algorithms could not process. As an example, on the PBMC data (∼ 68*k*), scImpute, drImpute and MAGIC exhausted the workstation memory. These algorithms were initially run on an 8GB RAM machine, observing that none of these algorithms ran successfully on this data, we tried running them on a system with an Intel Xeon 2.8 GHz processor having 64 GB of RAM. Still, none of these three imputation strategies were able to successfully impute this large an expression data. Our technique, efficiently ran and proved to be beneficial for this huge dataset.

### 3.2 Improvement in clustering accuracy

Clustering single cell RNA-seq data for discovering distinct cell types from a heterogeneous cell population is one of the most important applications of scRNA-seq. But, an algorithm which aims to cluster cells of similar types might get tricked by a large number of dropouts in single cell RNA seq data which serve as biological noise in the input to clustering algorithm. This incorrect view of expression levels should be fixed by a reasonable imputation resulting in accurate delineation of cell types. Hence, we observe the K-means clustering results on all the log-transformed expression profiles for each dataset both without and with imputation. The initialization parameter *K* (no of clusters) in K-means algorithm has been set to the number of annotated cell types for every data. Adjusted Rand Index (ARI) was used as the performance metric to evaluate the correspondence between the original annotations and K-means assigned clusters. Figure 2 clearly shows not only that 1 layer matrix factorization based expression re-estimation is the most beneficial as compared to other methods, but also, as we go deeper to 2 and 3 layers, we observe better performance for all datasets.

**Figure 2:**
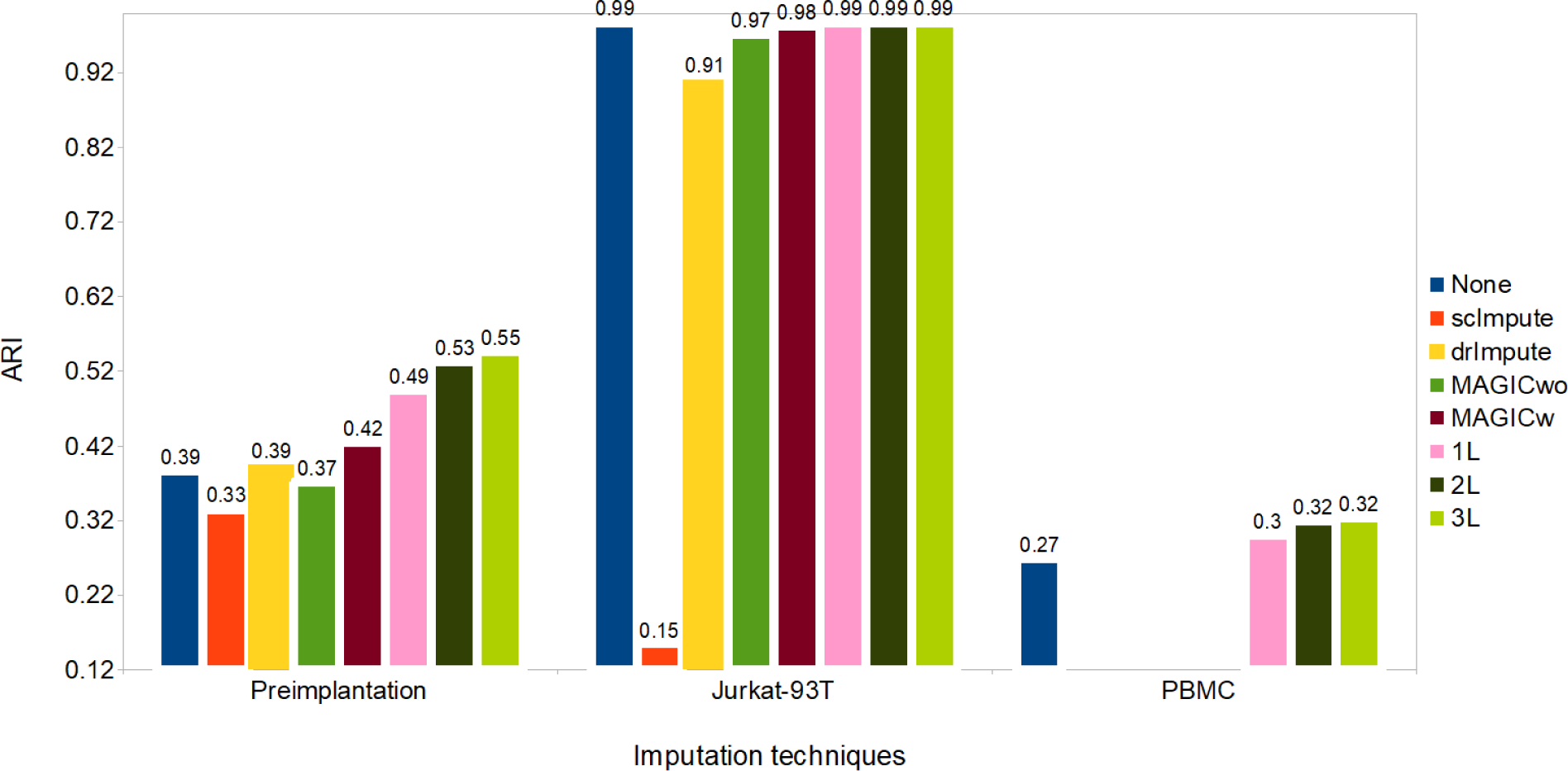
ARI values (rounded upto 2 decimal places) obtained after applying k-means clustering post various imputation techniques.

### 3.3 Improved differential Genes prediction

A good imputation method should result in better congruence between scRNA-seq and bulk RNA-seq data of the same biological condition on differentially expressed genes [10].

To assess the accuracy of differential expression (DE) analysis, We used the standard Wilcoxon Rank-Sum test for identifying differentially expressed genes from matrices obtained from various imputation methods. The dataset of pluripotent stem cells [32] (having matching bulk RNA-Seq data) generated from different individuals was used. The authors call it Tung dataset. They identify DE and non-DE genes using three standard methods: limma-voom [33, 34], edgeR [35] and DESeq2 [36] ^1^.

We show the agreement between bulk and single cell based DE calls using the Area Under the Curve (AUC) values obtained from the Receiver Operating Characteristic (ROC) curve (figure 3). The 2-layer (2L) deepMc imputation shows the best performance in predicting differentially expressed genes.

**Figure 3:**
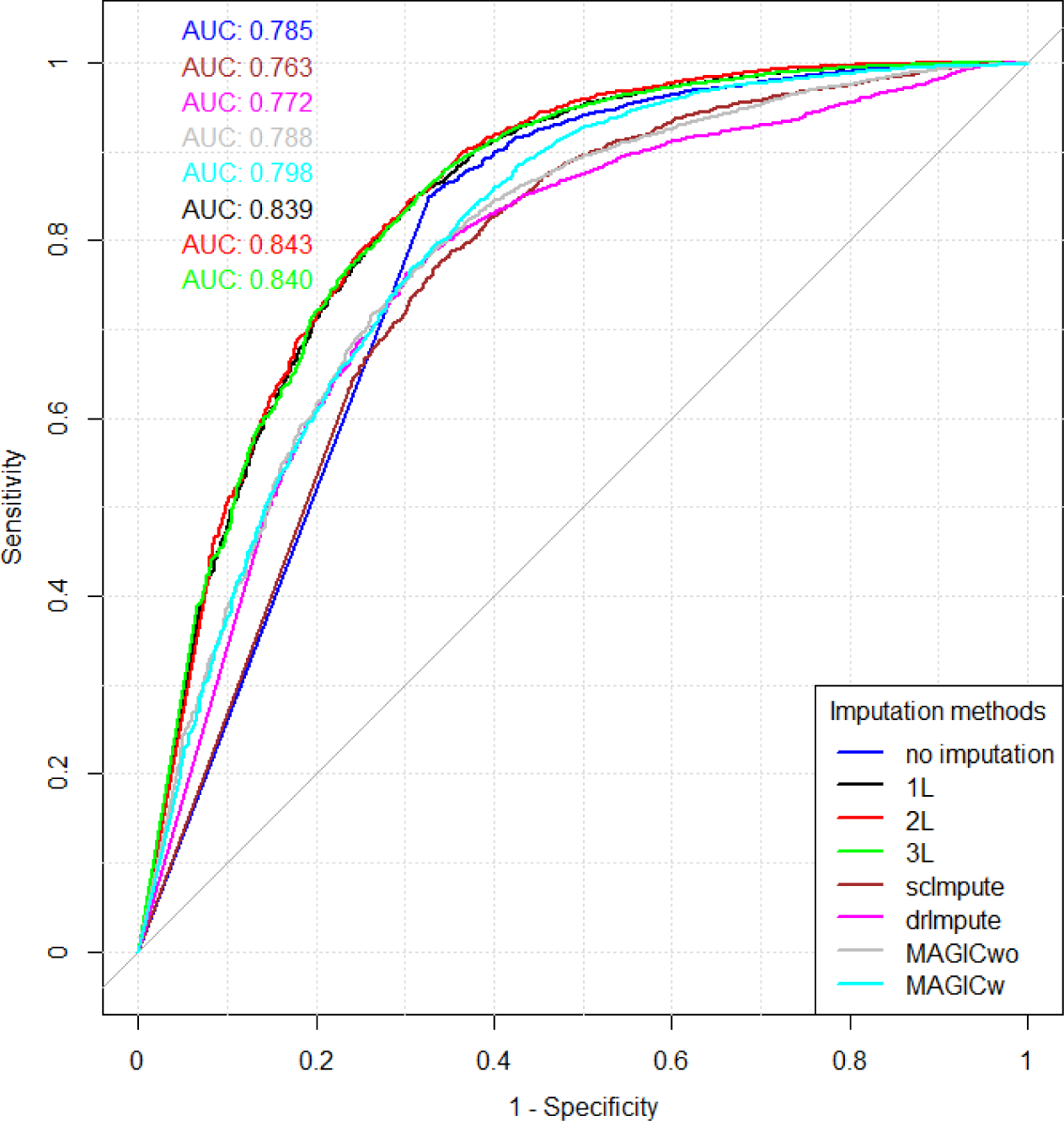
Plot showing ROC curve and AUC depicting how well do the DE genes predicted from scRNA and matching bulk RNA-Seq data agree. DE calls were made on expression matrix imputed using various methods.

For each method, the AUC value was computed on the identical set of ground truth genes. We had to make an exception only for drImpute as it applies an additional filter to prune genes. Hence, the AUC value was computed based on a smaller set of ground truth genes for drImpute.

### 3.4 Improvement in cell type separability

Before explaining this analysis, we introduce the following terms:

1. **Intra-cell type scatter**: For any two cell groups, we first find the median of Pearson’s correlation values computed for each possible pair of cells within their respective groups. We define the average of the median correlation values as the intra-cell type scatter.
2. **Inter-cell type scatter**: is defined as the median of Pearson’s correlation values computed for pairs such that in each pair, cells belong to two different groups.
3. **Cell-type separability (CTS) score:** The difference between the intra-cell scatter and inter-cell type scatter is termed as the cell-type separability (CTS) score.

An effective imputation should lead to a higher CTS score, showing that expression similarities between cells of identical type are considerably higher than that of cells coming from different subpopulations. The table S1 shows the resulting CTS before imputation and after applying various imputation strategies on any two cell types selected from each dataset. DeepImpute proves to be highly beneficial for 3 out of 4 datasets, with 2-layer (2L) and 3-layer (3L) deepMc algorithms giving the highest CTS score, validating our imputation strategy. We show the variation of CTS with various imputation strategies for Preimplantation dataset in figure 4 (others have been relegated to supplementary) and observe that our algorithms best separates the cell types, giving the highest CTS.

**Figure 4:**
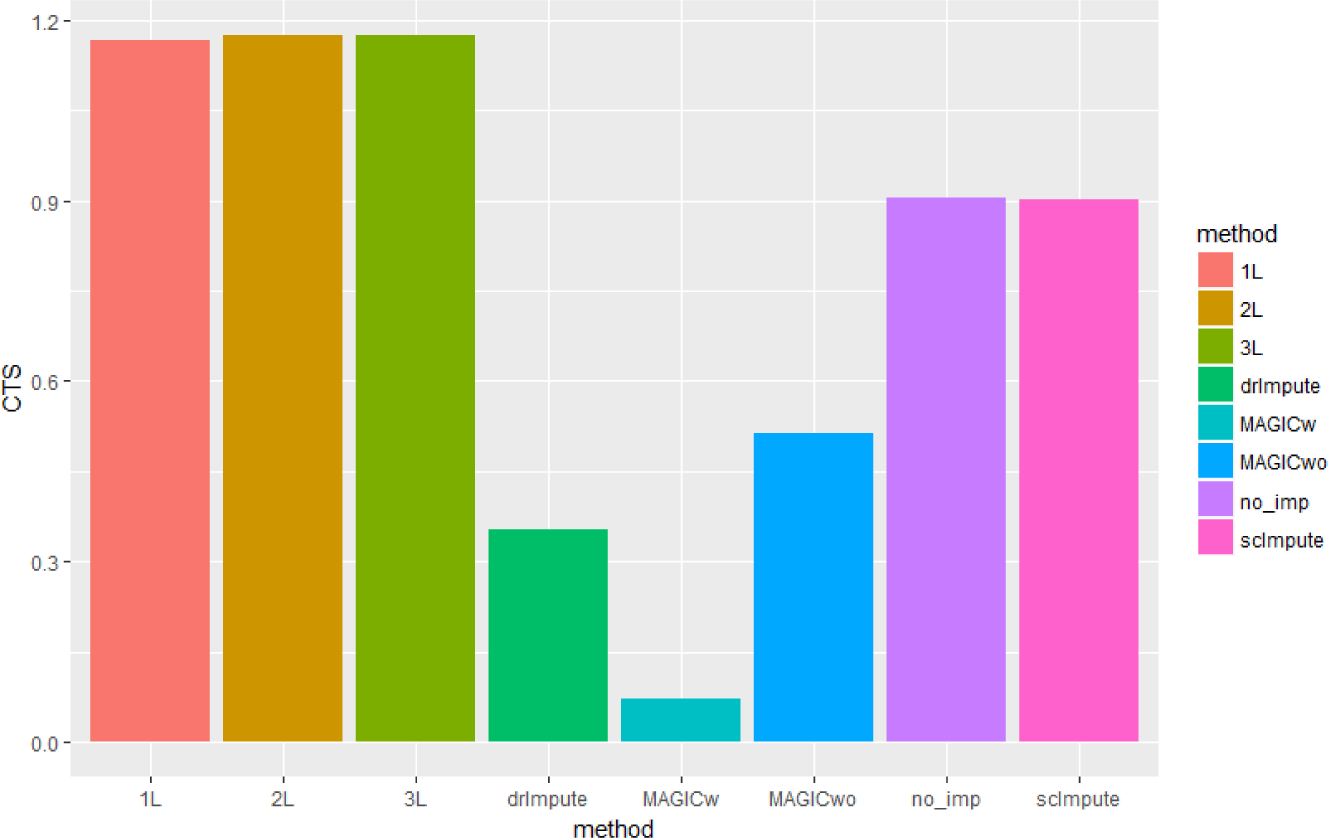
Plot showing the variation of CTS with various imputation strategies for Preimplantation dataset. (Refer table S1 for exact numeric values)

## 4 Conclusion

In this work, we present and compare the potency of deepImpute through rigorous experimentations including clustering accuracy, differential genes prediction and cell type separability, validating biologically relevant and best gene expression recovery outperforming various state-of-the-art imputation methods. We firmly believe it to be the most efficient and scalable method (capable of imputing huge datasets, unlike other algorithms) so far for imputation of scRNA-seq data.

The introduction and success of deepMc, paves way for more such deep learning based techniques in the field of computational biology, specifically, gene expression recovery.

## Footnotes

1 https://github.com/hemberg-lab/scRNA.seq.course

## References

[1] Wagner, A., Regev, A. & Yosef, N. Revealing the vectors of cellular identity with single-cell genomics. Nature biotechnology 34, 1145–1160 (2016).

[2] Kolodziejczyk, A., Kim, J. K., Svensson, V., Marioni, J. & Teichmann, S. The technology and biology of single-cell rna sequencing. Molecular Cell 58, 610 – 620 (2015). URL http://www.sciencedirect.com/science/article/pii/S1097276515002610.

[3] Wang, Z., Gerstein, M. & Snyder, M. Rna-seq: a revolutionary tool for transcriptomics. Nature reviews genetics 10, 57–63 (2009).

[4] Rizzetto, S. et al. Impact of sequencing depth and read length on single cell rna sequencing data of t cells. Scientific Reports 7, 12781 (2017).

[5] Patel, A. P. et al. Single-cell rna-seq highlights intratumoral heterogeneity in primary glioblastoma. Science 344, 1396–1401 (2014).

[6] Tirosh, I. et al. Dissecting the multicellular ecosystem of metastatic melanoma by single-cell rna-seq. Science 352, 189–196 (2016).

[7] Yan, L. et al. Single-cell rna-seq profiling of human preimplantation embryos and embryonic stem cells. Nature structural & molecular biology 20, 1131–1139 (2013).

[8] Tang, F. et al. Tracing the derivation of embryonic stem cells from the inner cell mass by single-cell rna-seq analysis. Cell stem cell 6, 468–478 (2010).

[9] Biase, F. H., Cao, X. & Zhong, S. Cell fate inclination within 2-cell and 4-cell mouse embryos revealed by single-cell rna sequencing. Genome research 24, 1787–1796 (2014).

[10] Kharchenko, P. V., Silberstein, L. & Scadden, D. T. Bayesian approach to single-cell differential expression analysis. Nature methods 11, 740–742 (2014).

[11] van Dijk, D. et al. Magic: A diffusion-based imputation method reveals gene-gene interactions in single-cell rna-sequencing data. BioRxiv 111591 (2017).

[12] Li, H. et al. Reference component analysis of single-cell transcriptomes elucidates cellular heterogeneity in human colorectal tumors. Nature Genetics (2017).

[13] Sengupta, D., Rayan, N. A., Lim, M., Lim, B. & Prabhakar, S. Fast, scalable and accurate differential expression analysis for single cells. bioRxiv 049734 (2016).

[14] Li, W. V. & Li, J. J. scimpute: accurate and robust imputation for single cell rna-seq data. bioRxiv 141598 (2017).

[15] Kwak, I.-Y., Gong, W., Koyano-Nakagawa, N. & Garry, D. Drimpute: Imputing dropout events in single cell rna sequencing data. bioRxiv 181479 (2017).

[16] Zheng, G. X. et al. Massively parallel digital transcriptional profiling of single cells. Nature communications 8, 14049 (2017).

[17] Wen, Z., Yin, W. & Zhang, Y. Solving a low-rank factorization model for matrix completion by a nonlinear successive over-relaxation algorithm. Mathematical Programming Computation 4, 333–361 (2012).

[18] Lee, D. D. & Seung, H. S. Learning the parts of objects by non-negative matrix factorization. Nature 401, 788 (1999).

[19] Olshausen, B. A. & Field, D. J. Emergence of simple-cell receptive field properties by learning a sparse code for natural images. Nature 381, 607 (1996).

[20] Candés, E. J. & Tao, T. The power of convex relaxation: Near-optimal matrix completion. IEEE Transactions on Information Theory 56, 2053–2080 (2010).

[21] Recht, B., Fazel, M. & Parrilo, P. A. Guaranteed minimum-rank solutions of linear matrix equations via nuclear norm minimization. SIAM review 52, 471–501 (2010).

[22] Kapur, A., Marwah, K. & Alterovitz, G. Gene expression prediction using low-rank matrix completion. BMC bioinformatics 17, 243 (2016).

[23] Min, S., Lee, B. & Yoon, S. Deep learning in bioinformatics. Briefings in bioinformatics 18, 851–869 (2017).

[24] Li, Z. & Tang, J. Weakly supervised deep matrix factorization for social image understanding. IEEE Transactions on Image Processing 26, 276–288 (2017).

[25] Trigeorgis, G., Bousmalis, K., Zafeiriou, S. & Schuller, B. W. A deep matrix factorization method for learning attribute representations. IEEE transactions on pattern analysis and machine intelligence 39, 417–429 (2017).

[26] Tariyal, S., Majumdar, A., Singh, R. & Vatsa, M. Deep dictionary learning. IEEE Access 4, 10096–10109 (2016).

[27] van Dijk, D. et al. Magic: A diffusion-based imputation method reveals gene-gene interactions in single-cell rna-sequencing data. BioRxiv 111591 (2017).

[28] Li, W. V. & Li, J. J. scimpute: accurate and robust imputation for single cell rna-seq data. bioRxiv 141598 (2017).

[29] Koren, Y., Bell, R. & Volinsky, C. Matrix factorization techniques for recommender systems. Computer 42, 30–37 (2009).

[30] Markovsky, I. Exact system identification with missing data. In Decision and Control (CDC), 2013 IEEE 52nd Annual Conference on, 151–155 (IEEE, 2013).

[31] Liao, B., Guo, C., Huang, L. & Wen, J. Matrix completion based direction-of-arrival estimation in nonuniform noise. In Digital Signal Processing (DSP), 2016 IEEE International Conference on, 66–69 (IEEE, 2016).

[32] Tung, P.-Y. et al. Batch effects and the effective design of single-cell gene expression studies. Scientific reports 7, 39921 (2017).

[33] Ritchie, M. E. et al. limma powers differential expression analyses for rna-sequencing and microarray studies. Nucleic acids research 43, e47–e47 (2015).

[34] Law, C. W., Chen, Y., Shi, W. & Smyth, G. K. voom: Precision weights unlock linear model analysis tools for rna-seq read counts. Genome biology 15, R29 (2014).

[35] Zhou, X., Lindsay, H. & Robinson, M. D. Robustly detecting differential expression in rna sequencing data using observation weights. Nucleic acids research 42, e91–e91 (2014).

[36] Love, M. I., Huber, W. & Anders, S. Moderated estimation of fold change and dispersion for rna-seq data with deseq2. Genome biology 15, 550 (2014).

